# The Genomic Stability at the Coding Regions of the Multidrug Transporter Gene *ABCB1*: Insights into the Development of Alternative Drug Resistance Mechanisms in Human Leukemia Cells

**DOI:** 10.1101/820639

**Authors:** Kevin G. Chen, George E. Duran, Mark J. Mogul, Yan C. Wang, Kevin L. Ross, Jean-Pierre Jaffrézou, Lyn M. Huff, Tito Fojo, Norman J. Lacayo, Branimir I. Sikic

**Affiliations:** Division of Oncology, Department of Medicine, Stanford University School of Medicine, Stanford, California 94305, USA; NIH Stem Cell Unit, National Institute of Neurological Disorders and Stroke, National Institutes of Health, Bethesda, Maryland, USA; Medicine Branch, National Cancer Institute, National Institutes of Health, Bethesda, Maryland, USA; Division of Pediatric Hematology-Oncology-Stem Cell Transplantation and Cancer Biology, Stanford University School of Medicine and Stanford Cancer Institute, Palo Alto, California, USA

**Author notes:** J.P.J. French National Centre for Scientific Research, Paris, France; K.G.C. NIH Stem Cell Unit, National Institute of Neurological Disorders and Stroke, National Institutes of Health, MD USA; K.L.R., Ross BioPharm Group, Rocky Point, NY, USA; M.J.M. Medical Affairs U.S., Servier Pharmaceuticals, Boston, MA, USA; T.F. Herbert Irving Comprehensive Cancer Center, Columbia University Medical Center/New York-Presbyterian Hospital, New York, NY, USA. **Correspondence**: Kevin G. Chen, M.D., Ph.D, NINDS, NIH, 9000 Rockville Pike, Building 10, Room 3B14, Bethesda, MD, 20892 USA.

**Keywords:** cancer, leukemia, multidrug resistance, ABCB1, P-glycoprotein, cyclosporine, CFTR, mutation

## Abstract

Despite considerable efforts in reversing clinical multidrug resistance (MDR), targeting the predominant multidrug transporter ABCB1/P-glycoprotein (P-gp) based on small molecule inhibitors has been hindered. This may be due to the emergence of alternative drug resistance mechanisms. However, the non-specific P-gp inhibitor cyclosporine (CsA) showed significant clinical benefits in patients with acute myeloid leukemia (AML), which likely represents the only proof-of-principle clinical trial using several generations of MDR inhibitors. Nevertheless, the mechanisms that underlie this successful MDR modulation by CsA are not elucidated because of the absence of CsA-relevant cellular models. In this study, we report the development of two erythroleukemia variants, RVC and RDC, which were derived by step-wise co-selection of K562/R7 drug-resistant leukemia cells with the etoposide-CsA and doxorubicin-CsA drug combinations, respectively. Interestingly, both RVC and RDC, which retained P-gp expression, showed altered MDR phenotypes that were resistant to cyclosporine modulation. The *ABCB1* coding regions were genetically stable even under long-term stringent drug selection. Genomically, *ABCB1* is likely the most stable ABC transporter gene when comparing with several ABC superfamily members (such as *ABCA1*, *ABCC1*, *CFTR*, and *ABCG2*). Our findings suggested that non-P-gp mechanisms were likely responsible for the resistance to CsA modulation in both RVC and RDC cells. Moreover, we found that CsA played a role in undermining the selection of highly drug-resistant cells via induction of low level and unstable drug resistance, thus shedding some light on the benefits of CsA in treating certain types of AML patients.

## INTRODUCTION

Multidrug resistance (MDR), a phenotype that has been well-defined both *in vitro* in cell culture and *in vivo* in cancer patients, is principally caused by numerous ATP-binding cassette (ABC) transporters [reviewed in (Robey et al., 2018)]. The best characterized ABC transporter is ABCB1, known as P-glycoprotein (P-gp), which was implicated in clinical MDR [reviewed in (Amiri-Kordestani et al., 2012; Chen and Sikic, 2012; Robey et al., 2018; Sikic et al., 1997; Tamaki et al., 2011)]. A wide range of clinical trials were conducted over the past 30 years in patients with different types of cancer by coadministration of the small molecule inhibitors of P-gp and anticancer drugs. There were three generations of P-gp inhibitors that were used in these trials, including cyclosporine (CsA) (Bartlett et al., 1994; List et al., 2001; List et al., 1993; Liu Yin et al., 2001; Lum et al., 1992; Yahanda et al., 1992), the cyclosporine D analogue PSC 833 (PSC) (Advani et al., 2001; Advani et al., 2005; Dorr et al., 2001; Fracasso et al., 2000), and zosuquidar (Cripe et al., 2010; Lancet et al., 2009; Marcelletti et al., 2019).

Despite these substantial clinical efforts, most of the advanced phase III trials showed no favorable outcomes using highly specific P-gp inhibitors (e.g., PSC and zosuquidar). Thus, it was postulated that there might be redundant drug-resistant mechanisms that are responsible for resistance to MDR modulation in cancer cells (Amiri-Kordestani et al., 2012; Chen and Sikic, 2012; Robey et al., 2018; Sikic et al., 1997; Tamaki et al., 2011). The mechanisms that underlie resistance to MDR modulation are complicated and multifactorial in nature. These underlying mechanisms comprise: (i) the genetic heterogeneity of the *ABCB1* gene (e.g., various single nucleotide polymorphisms and rare mutations) (Chen et al., 1997; Chen et al., 2000; Choi et al., 1988; Kimchi-Sarfaty et al., 2007; Morisaki et al., 2005; Zu et al., 2014); (ii) genomic and epigenomic instability events that regulate *ABCB1* expression (Chen et al., 1994; Chen et al., 2005; Huff et al., 2005; Mickley et al., 1998; Patch et al., 2015); (iii) the involvement of non-P-gp ABC transporters (e.g., *ABCA3*, *ABCC1/MRP1*, and *ABCG2/BCRP*) (Calcagno et al., 2008; Cole et al., 1992; Doyle et al., 1998; Steinbach et al., 2006; van der Kolk et al., 2002; Wulf et al., 2004; Zhan et al., 1997); and (iv) various non-MDR modifiers that involve cell cycle control, anti-apoptosis, and cytoskeleton alterations [reviewed in (Chen and Sikic, 2012; Dumontet and Sikic, 1999; Giannakakou et al., 2000)].

Nevertheless, a significant clinical benefit, achieved in elderly refractory AML patients in a trial using the non-specific P-gp inhibitor CsA (List et al., 2001), is overshadowed due to the results of subsequent CsA trials under different conditions (Liu Yin et al., 2001), perhaps resulting from a flawed trial design (Smith et al., 2003) and the selection of inappropriate AML subtypes. Both potential factors made CsA data interpretation quite difficult and with inconclusive results.

CsA-mediated modulation of MDR still represents the only proof-of-principle clinical trial of MDR modulation. Moreover, CsA-mediated modulation of MDR is poorly understood, largely owing to the absence of drug-resistant models based on CsA-mediated P-gp inhibition. Several existing MDR modulation models were based on the use of the potent and specific P-gp inhibitor PSC (Beketic-Oreskovic et al., 1995; Chen et al., 1997; Dumontet et al., 2004; Duran et al., 2015; Jaffrezou et al., 1995), which provided insights into alternative drug-resistance (ADR) mechanisms that are not relevant to CsA modulation.

In this study, we report the establishment of CsA-relevant models, RVC and RDC, by step-wise selection of K562/R7 P-gp-overexpressing leukemia cells with the etoposide-CsA and doxorubicin-CsA combinations, respectively. We characterized their phenotypes based on drug-resistant profile and drug accumulation. We also examined the mutational events of *ABCB1* in these cell lines, with a focus on drug-induced or drug-selected mutations at the mRNA coding region. In addition, we compared the variant or mutational spectrum of several representative transporters from the ABC superfamily to assess the involvement of gene mutations in the resistance to MDR modulation. Lastly, we proposed several relevant models that account for the CsA effects on deferring the onset of stable and highly drug-resistant phenotypes in leukemia cells.

## RESULTS

### Destabilizing the stable multidrug resistant (MDR) phenotypes in K562 cells by cyclosporine

We selected drug-resistant leukemic cell lines by step-wise co-selection of non-P-gp expressing erythroleukemic cell line K562 with tubulin active drugs paclitaxel (Taxol) and vinblastine in the presence of 2 μM CsA (Figure 1A, 1B). However, following a one-year period of drug selection, we were unable to derive stable drug-resistant lines. Typically, the selected cells lost drug resistance to the selecting agents following drug-free growth conditions for 2 to 3 weeks. The drug-resistant phenotypes were stabilized when CsA was replaced with PSC in the selection regimens. The resulting cell lines KPTA5 and KCVB2 cells exhibited non-ABC transporter mechanisms that are associated with altered tubulin expression and polymerization (Dumontet et al., 2004; Jaffrezou et al., 1995). Nonetheless, the results from CsA-selection regimens suggested that CsA may play a role in delaying the emergence of stable drug-resistant phenotypes.

**Figure 1.**
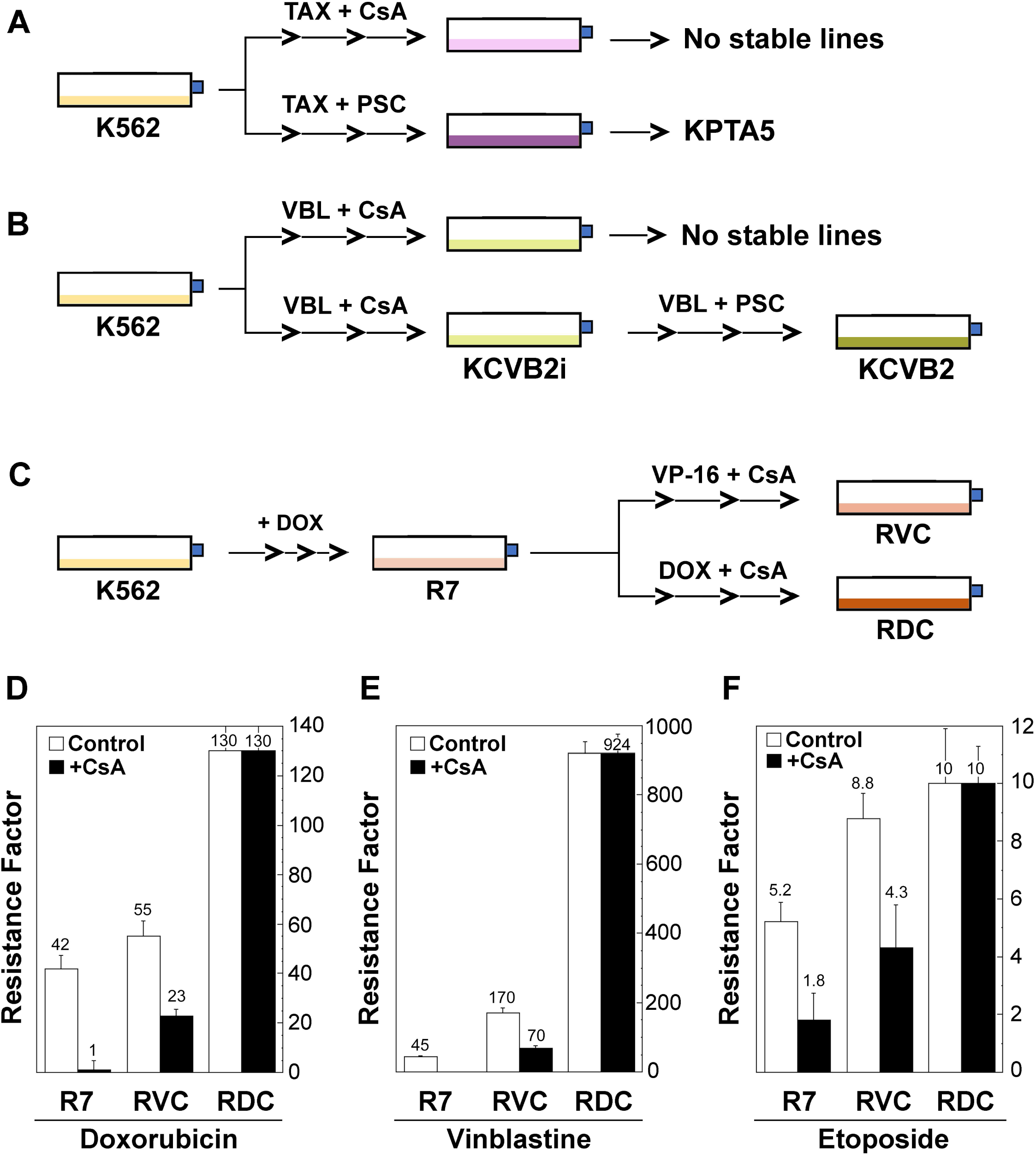
Development of drug-resistant leukemic cell lines that are insensitive to MDR modulation. (**A** and **B**) Development of the paclitaxel (taxol)- and vinblastineresistant leukemic cell lines (KPTA5 and KCVB2) in the presence of cyclosporine (CsA) or PSC 833 (PSC). (**C**) Development of drug-resistant leukemic cell lines (RVC and RDC) by step-wise co-selection of the multidrug-resistant line K562/R7 (R7) with etoposide (VP-16) or doxorubicin (DOX) in the presence of 2 μM cyclosporine (CsA). (**D**) Drug resistance factors represent fold changes of drug resistance relative to the parental K562 cells as determined by the MTT assays. The mean values, labeled on the top of the columns, of resistance factors were shown from one of the two similar experiments.

### Development of MDR leukemic lines that are insensitive to cyclosporine modulation in K562/R7 cells

To derive stable drug-resistant lines that were insensitive to CsA modulation, we coselected the P-gp-expressing line K562/R7 (R7) with exposure of the cells to etoposide (VP-16) and doxorubicin in the presence of 2 μM CsA. The derived lines were named RVC and RDC, respectively (Table 1, Figure 1C). These two cell lines expressed high levels of *ABCB1* mRNA by RT-PCR and high levels of P-gp like the parental R7 by Western blotting using the monoclonal antibody C219.

**Table 1.**
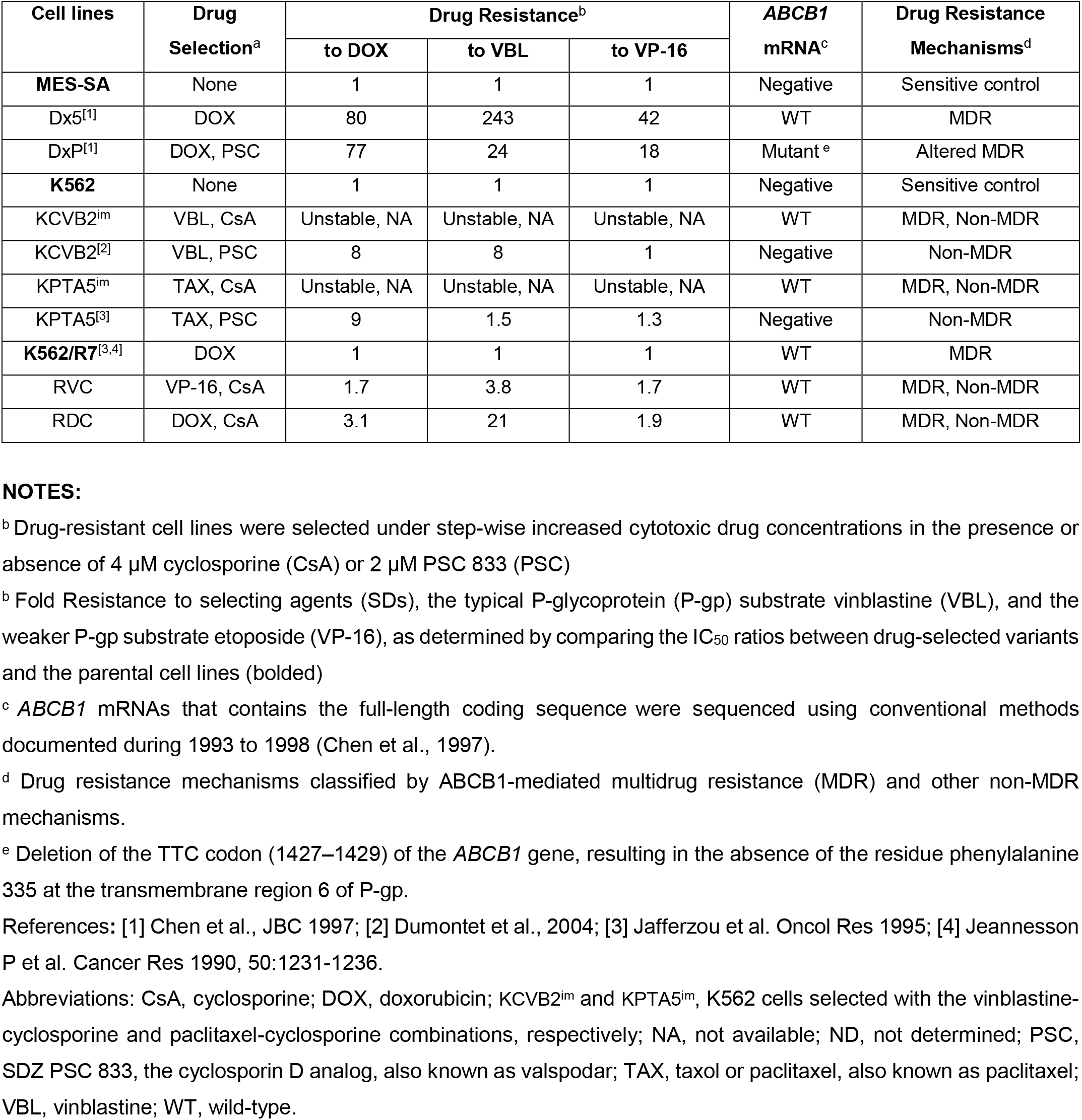
Development of Multidrug Resistant Cell Lines that are Insensitive to Cyclosporine Modulation

| Cell lines | Drug Selection<sup>a</sup> | Drug Resistance<sup>b</sup> |  |  | *ABCB1* mRNA<sup>c</sup> | Drug Resistance Mechanisms<sup>d</sup> |
| --- | --- | --- | --- | --- | --- | --- |
|  |  | to DOX | to VBL | to VP-16 |  |  |
| **MES-SA** | None | 1 | 1 | 1 | Negative | Sensitive control |
| Dx5<sup>[1]</sup> | DOX | 80 | 243 | 42 | WT | MDR |
| DxP<sup>[1]</sup> | DOX, PSC | 77 | 24 | 18 | Mutant<sup>e</sup> | Altered MDR |
| **K562** | None | 1 | 1 | 1 | Negative | Sensitive control |
| KCVB2<sup>im</sup> | VBL, CsA | Unstable, NA | Unstable, NA | Unstable, NA | WT | MDR, Non-MDR |
| KCVB2<sup>[2]</sup> | VBL, PSC | 8 | 8 | 1 | Negative | Non-MDR |
| KPTA5<sup>im</sup> | TAX, CsA | Unstable, NA | Unstable, NA | Unstable, NA | WT | MDR, Non-MDR |
| KPTA5<sup>[3]</sup> | TAX, PSC | 9 | 1.5 | 1.3 | Negative | Non-MDR |
| **K562/R7**<sup>[3,4]</sup> | DOX | 1 | 1 | 1 | WT | MDR |
| RVC | VP-16, CsA | 1.7 | 3.8 | 1.7 | WT | MDR, Non-MDR |
| RDC | DOX, CsA | 3.1 | 21 | 1.9 | WT | MDR, Non-MDR |

**NOTES:**
<sup>b</sup> Drug-resistant cell lines were selected under step-wise increased cytotoxic drug concentrations in the presence or absence of 4 μM cyclosporine (CsA) or 2 μM PSC 833 (PSC)
<sup>b</sup> Fold Resistance to selecting agents (SDs), the typical P-glycoprotein (P-gp) substrate vinblastine (VBL), and the weaker P-gp substrate etoposide (VP-16), as determined by comparing the IC<sub>50</sub> ratios between drug-selected variants and the parental cell lines (bolded)
<sup>c</sup> *ABCB1* mRNAs that contains the full-length coding sequence were sequenced using conventional methods documented during 1993 to 1998 (Chen et al., 1997).
<sup>d</sup> Drug resistance mechanisms classified by *ABCB1*-mediated multidrug resistance (MDR) and other non-MDR mechanisms.
<sup>e</sup> Deletion of the TTC codon (1427–1429) of the *ABCB1* gene, resulting in the absence of the residue phenylalanine 335 at the transmembrane region 6 of P-gp.
References: [1] Chen et al., JBC 1997; [2] Dumontet et al., 2004; [3] Jafferzou et al. Oncol Res 1995; [4] Jeannesson P et al. Cancer Res 1990, 50:1231-1236.
Abbreviations: CsA, cyclosporine; DOX, doxorubicin; KCVB2<sup>im</sup> and KPTA5<sup>im</sup>, K562 cells selected with the vinblastine-cyclosporine and paclitaxel-cyclosporine combinations, respectively; NA, not available; ND, not determined; PSC, SDZ PSC 833, the cyclosporin D analog, also known as valsopodar; TAX, taxol or paclitaxel, also known as paclitaxel; VBL, vinblastine; WT, wild-type.

Both RVC and RDC cells were moderately resistant (1.7- and 3.1-fold) to their selecting drugs, etoposide and doxorubicin respectively, but highly resistant (21-fold) to the typical P-gp substrate vinblastine in RDC cells (Figure 1D-F). In contrast to R7 cells, whose resistance to doxorubicin was significantly modulated by CsA (97.6%) (Figure 1D, columns 1 and 2), we observed partial modulation of doxorubicin resistance (58%) under identical experimental conditions (Figure 1D, columns 3 and 4). In particular, RDC cells exhibited complete resistance to the modulatory effect of CsA (Figure 1D: columns 5 and 6).

With respect to the capacity of CsA to modulate vinblastine resistance, we observed a complete reversion of vinblastine resistance (45-fold) to the level of parental K562 cells in R7 cells, but partial revision (~59% of reversion) in RVC and insensitive in RDC cells (Figure 2E). Concerning the ability of CsA to reverse etoposide resistance, we observed 65% of reversion of the resistance in R7 cells, 51% in RVC, and no modulation in RDC cells (Figure 2F). These data suggested the co-selected cell lines RVC and RDC had altered MDR phenotypes that were insensitive to the MDR inhibitor CsA.

**Figure 2.**
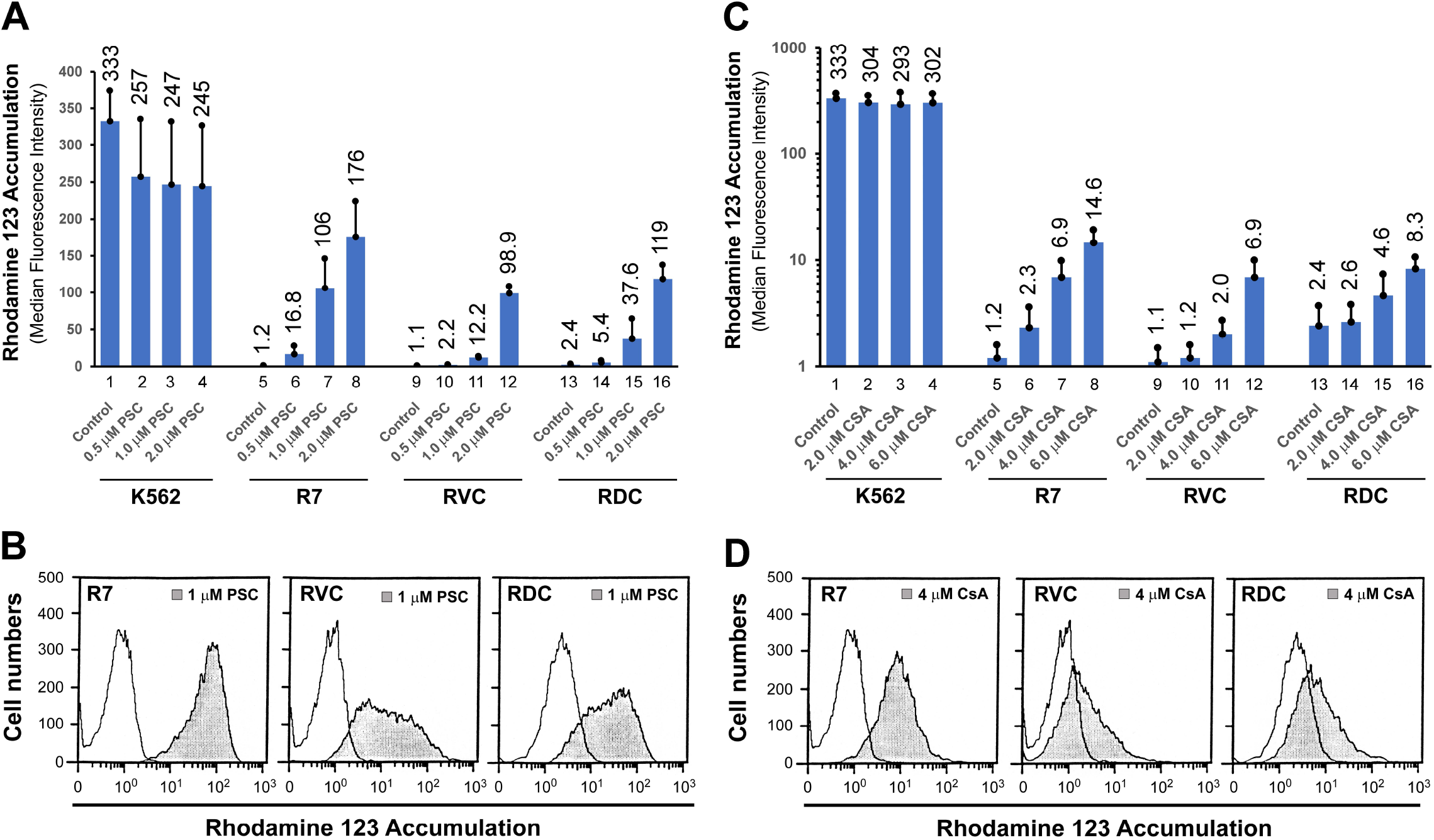
Rhodamine 123 (Rh-123) accumulation assays. (**A and B**) Rh-123 accumulation in the presence or absence of cyclosporine (CsA) and PSC 833 (PSC) in K562, R7, RVC, and RDC cells. The mean values (columns), also labeled on the top of the histograms, and standard deviations (bars) are derived from quadruplicate determinants. (**C and D**) Representative histograms of Rh-123 accumulation in R7, RVC, and RDC cells in the presence of 1 μM PSC and 4 μM CsA.

### Fluorescent dye Rhodamine 123 (Rh-123) accumulation

Rh-123 is a typical P-gp substrate, whose accumulation can be easily detected by flow cytometry. In P-gp negative K562 cells, Rh-123 accumulation, served as 100% control, was reduced by the potent P-gp inhibitor PSC (Figure 2A, columns 1-4) and was insensitive to the effects of various CsA concentrations (Figure 2C, columns 1-4). Of note, the mechanisms underlying the effect of PSC on decreased Rh-123 accumulation in Pgp negative cells are unclear.

In P-gp-expressing R7 cells, Rh-123 accumulation was 0.4% of the K562 control, which was reversed by PSC in a dose-dependent manner and up to 53% of the K562 control by 2 μM PSC (Figure 2A, columns 1, 5 to 8). The Rh-123 defect in both RVC and RDC cell lines was less sensitive to 2 μM PSC, displaying 30% and 36% of reversion, respectively, under these identical conditions (Figure 2A, columns 1, 12, and 16).

Moreover, Rh-123 accumulation in R7 cells (i.e., 0.4% of K562 control) was reversed to 0.7%, 2.0%, and 4.4% of the K562 control level by 2, 4, and 6 μM CsA (Figure 2C, columns 5-8). Rh-123 transport in both RVC and RDC cells was less sensitive to CsA modulation under these identical conditions. For example, under 4 μM CsA, a 5.8-fold increase in Rh-123 accumulation was found in R7 cells (Figure 2C, columns 5 and 7), but, only a 1.8- and 1.9-fold increase in Rh-123 signal in RVC and RDC cells, respectively (Figures 2C: columns 9 and 11, 13 and 15; Figure 2D). Taken together, the Rh-123 accumulation patterns in both RVC and RDC cells support an altered MDR phenotype that may be associated with defective drug transport mechanisms in these cell lines.

### Anticancer drug accumulation in RVC and RDC lines

To further define the altered drug transport phenotypes, we evaluated the intracellular accumulation of [^3^H]-labeled drugs. As shown in Figure 3A and Table 2, all R7, RVC, and RDC lines showed reduced [^3^H]-daunorubicin compared to both K562 and R7. The [^3^H]-daunorubicin accumulation was increased 2.9-, 3.0-, and 2.3-fold in R7 cells treated with CsA, PSC, and verapamil, respectively. However, [^3^H]-daunorubicin accumulation was only increased by 2.3-, 2.3-, and 1.7-fold in RVC cells, and 2.1-, 2.2-, and 1.6-fold in RDC cells under identical conditions (Figure 3A).

**Figure 3.**
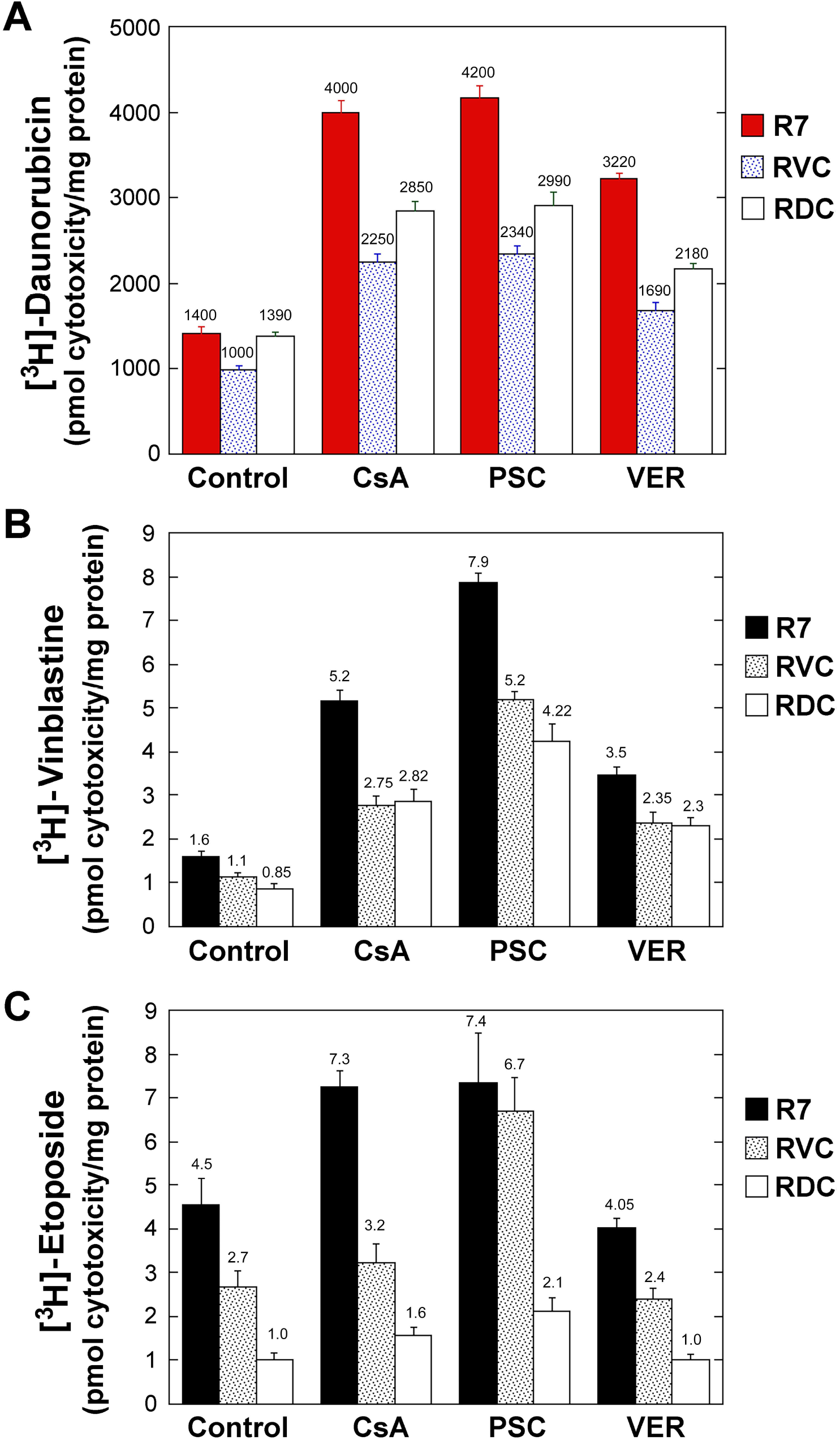
Intracellular cytotoxic drug accumulation. Measurements of the intracellular [^3^H]-daunorubicin (**A**), [3H]-vinblastine (**B**), and [3H]-etoposide (**C**) concentrations in the presence or absence of cyclosporine (CsA), PSC 833 (PSC), and verapamil (VER). The mean values (columns), also labeled on the top of the histograms, and standard deviations (bars) are derived from quadruplicate determinants. One of two similar experiments is shown.

**Table 2.** Comparative Analysis of Drug Export in Multidrug Resistant Cell Lines

| **Drugs** | **R7** | **RVC** | **RDC** |
| --- | --- | --- | --- |
| [<sup>3</sup>H]-Daunorubicin | 1* | 1* | 1* |
| + CsA | 2.9 | 2.3 | 2.1 |
| + PSC | 3.0 | 2.3 | 2.2 |
| + VER | 2.3 | 1.7 | 1.6 |
| [<sup>3</sup>H]-Vinblastine | 1* | 1* | 1* |
| + CsA | 3.3 | 2.5 | 3.3 |
| + PSC | 4.9 | 4.7 | 5.0 |
| + VER | 2.2 | 2.1 | 2.7 |
| [<sup>3</sup>H]-Etoposide | 1* | 1* | 1* |
| + CsA | 1.6 | 1.2 | 1.6 |
| + PSC | 1.6 | 2.5 | 2.1 |
| + VER | 0.9 | 0.9 | 1.0 |

**NOTES:**

Moreover, the defect of [^3^H]-vinblastine accumulation was increased 3.3-, 4.9-, and 2.2-fold in R7 cells treated with CsA, PSC, and verapamil respectively (Figure 3B, Table 2). However, there were similar [^3^H]-vinblastine accumulation patterns in both RVC and RDC cells, displaying 2.5-, 4.7-, and 2.1-fold in RVC cells, 3.3-, 5.0-, and 2.7-fold increases in RDC cells under these identical modulation conditions (Figure 3B, Table 2). Thus, the [^3^H]-vinblastine accumulation patterns suggest the coexistence of a functional P-gp status in both RVC and RDC cells.

In addition, [^3^H]-etoposide uptake was significantly decreased in both RVC and RDC, exhibiting a 1.7-fold and 4.5-fold reduction in drug accumulation in RVC and RDC cells, respectively, as compared to R7 cells (Figure 3C, columns 1-3). Likewise, this defective pattern was unable to be reversed to the level of parental R7 cells when RVC and RDC cells were treated with the non-specific P-gp inhibitor CsA, rather than the specific P-gp inhibitor PSC (Figure 3C). There was a 1.6-, 1.6-, and 0.9-fold increase in [^3^H]-etoposide in R7 cells by CsA, PSC, and verapamil treatments respectively. Still, there was a similar modulation pattern in both RVC and RDC cells compared with R7 cells, displaying 1.2-, 2.5-, and 0.9-fold increase in RVC cells; 1.6-, 2.1-, and 1.0-fold increase in RDC cells, in the presence of CsA, PSC, verapamil, respectively (Figure 3B, Table 2). These data indicated that the defect of [^3^H]-etoposide was not mainly mediated by P-gp, suggesting that mutant P-gp or non-P-gp transporters may be responsible for this phenotype.

Taken together, [^3^H]-labeled drug uptake together with Rh-123 retention revealed drug accumulation defects in both RVC and RDC cells, which was consistent with the evolution of a mutant P-gp, or post-translationally modified P-gp, or non-P-gp drug resistance mechanisms that rendered cells resistant to both the selecting drugs (i.e., doxorubicin and etoposide) and the MDR modulator cyclosporine. The decreased drug accumulation in these cells are also consistent with an alternate transporter mechanism.

### Phe^335^ (F335) codon in R7, RVC, RDC, and human cancers

We previously identified the F335 (coded by the 1247-1249 TTC) deletion in P-gp in a mutant MDR line (DxP), derived from MES-SA/Dx5 cells by step-wise co-selection of the cells with doxorubicin and PSC (Chen et al., 1997; Chen et al., 2000). This deletion led to DxP cells resistant to the reversing effects of MDR inhibitors such as CsA and PSC 833 (Chen et al., 1997). Thus, we further examined the frequency of this mutation in R7, RVC, and RDC cells, as well as, in 24 *ABCB1*-expressing human cancer samples. These samples included 9 leukemias, 6 lymphomas, and 6 solid malignant tumors. Using mutant *ABCB1* PCR products from DxP cells as the template, we performed RT-PCR and cDNA duplex assays. However, the assays failed to detect the 1247-1249 TTC deletion in all 24 cancer samples, which suggested that the F335 deletion might be an uncommon event for human *ABCB1* in drug-selected cancer cell lines and in human cancer patients.

### *ABCB1* sequencing analysis in R7, RVC, RDC, and KCVB2 cells

We manually sequenced the whole coding regions of the *ABCB1* gene in R7, RVC, RDC, and KCVB2i. KCVB2i, is an intermediate line of KCVB2, which had a low level of P-gp expression (Table 1). We found no mutations in the coding regions of *ABCB1* in R7, RVC, RDC, and KCVB2i cells (Table 1). These data suggested that the *ABCB1* coding regions are stable and resistant to drug-induced mutations in human leukemia cells. Thus, our data ruled out the possibility that the altered MDR phenotypes in both RVC and RDC were due to the contribution of a mutant P-gp.

### Upstream *ABCB1* activation in leukemic MDR cell lines

To examine the role of *ABCB1* regulation in drug-resistant RVC and RDC cells, we used an RNase protection assay to determine *ABCB1* mRNA expression and to map the transcription start sites of the *ABCB1* gene. As indicated in Figure 4, RNase assay identified two protected fragments: one at exon 1b and another one at the upstream regions of exon 1a. Although the exact upstream starting site(s) could not be identified in this assay, the data indicated that *ABCB1* expression in these drug-resistant variants was regulated by two different transcriptional mechanisms. Interestingly, etoposide-CsA coselected RVC cells showed a similar RNase protection pattern to that of parental R7 cells, which suggested that this co-selection regimen might have less impact on *ABCB1* mRNA and protein expression. But, doxorubicin-CsA co-selected RDC cells showed a more than 50% decrease in upstream transcripts concomitantly with an increase in *ABCB1* mRNA expression initiated from exon 1b. Thus, RDC data indicated a positive selection of the native *ABCB1* promoter (P1) for the regulation of *ABCB1* and P-gp expression in these cells. The lack of *ABCB1* mRNA mutations without significantly altering *ABCB1* upstream transcription prompted us to compare the variations or mutations among diverse ABC drug transporters

**Figure 4.**
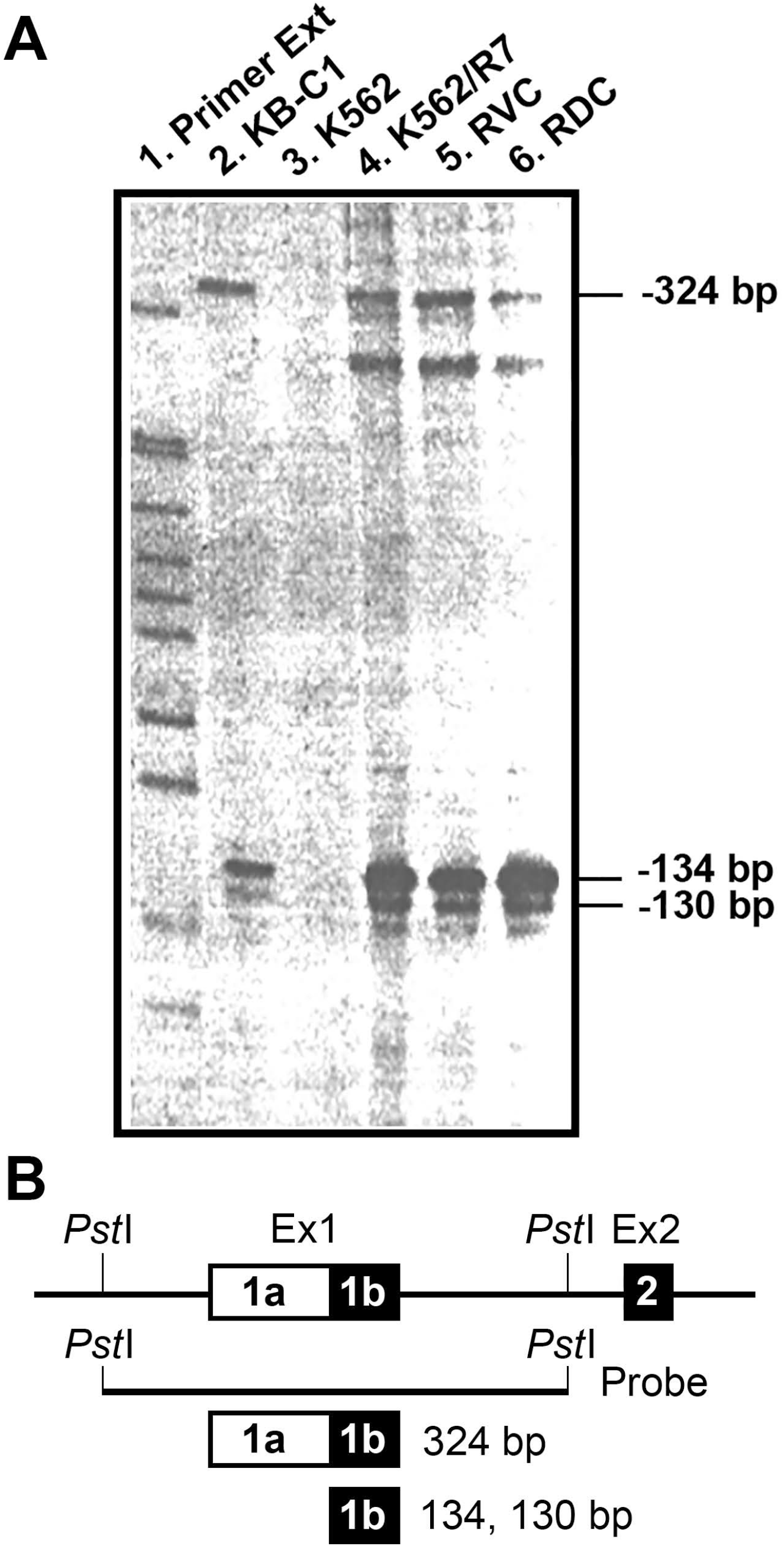
RNase protection analysis of upstream transcriptional start sites. (**A**) Total RNAs, extracted from K562 parental cells (*ABCB1* mRNA negative control), KB-C1 cells (*ABCB1* mRNA positive control), and three drug-resistant variants (K562/R7, RVC, RDC), were hybridized with the antisense RNA probe described below in Figure 4B. Primer extension products (lane 1) were used to map the locations of the transcription starting sites (−324-bp, and −130/-134-bp) upstream of the translation starting codon of the *ABCB1* gene. (**B**) Structural presentation of the genomic probe (the 982-bp *Pst*I-*Pst*I genomic fragment) of the *ABCB1* proximal promoter used for generating the antisense probe used in the RNase protection assays

### Genetic stability and instability of ABC transporters at the coding regions

There are 48 ABC transporters in the human genome. We analyzed the genomic stability in several representative ABC transporters (e.g., *ABCB1*, *ABCC1*, *ABCG2*, *ABCA1*, and *ABCC7*/*CFTR*) based on the curated UniProt database (www.uniprot.org) (Figure 5). ABCB1, ABCC1, and ABCG2 are well-studied multidrug transporters *in vitro*. ABCA1 functions as an exporter for intracellular cholesterol and certain phospholipids. CFTR, possibly the only ABC transporter that has gating channel activity, functions as a chloride and bicarbonate exporter (www.uniprot.org).

**Figure 5.**
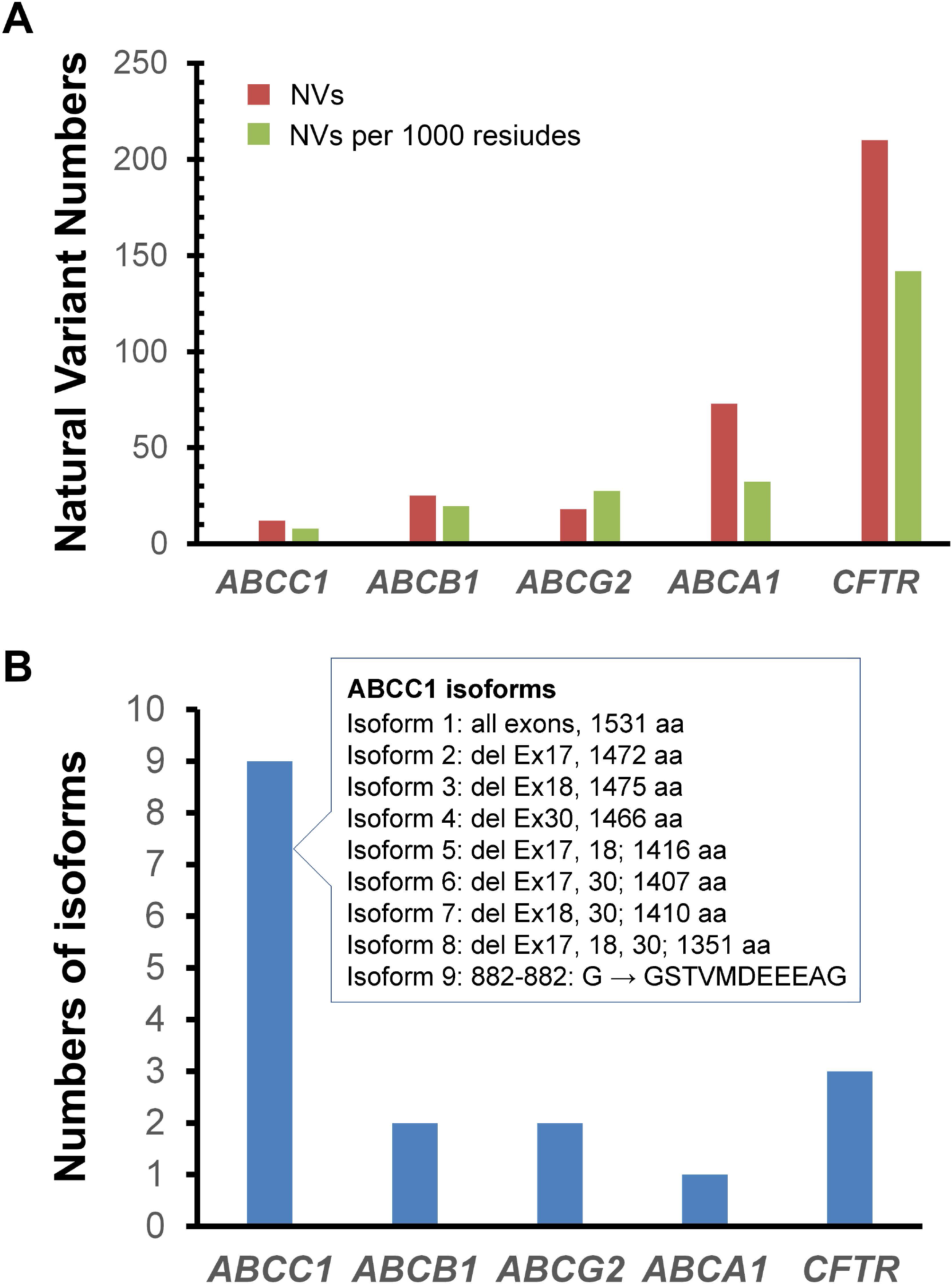
Coding sequence analysis of representative ABC transporters. **(A)** Frequency of natural variants in several representative ABC transporters. **(B)** Isoforms of the representative ABC transporters. All data were retrieved from the UniProt database (www.uniprot.org) as of December 2018. Abbreviations: aa, amino acid residues; del, deletion; Ex, exon; NVs, natural variants.

We analyzed natural variants, including polymorphisms, variations, and disease-associated mutations at the coding regions of these transporter genes. As shown in Figure 5A, *ABCC1*, *ABCB1*, *ABCG2*, *ABCA1*, and *CFTR* had the frequencies of 8, 20, 28, 32, and 142 natural variants per 1000 residues, respectively (Figure 5A). Thus, *ABCC1* appeared to be the most stable gene in terms of its frequency of natural variants in the coding regions in these analyses. (Figure 5A). Furthermore, *ABCG2* and *ABCA1* had a 1.4- and 1.6-fold increase in the frequency of natural variants relative to *ABCB1*. Surprisingly, the natural variant frequency of CFTR was 7.3-fold higher than that of *ABCB1* (Figure 5A). Regardless of these natural variants, *ABCC1* had 9 isoforms, with deletions frequently occurring at exon 17, 18, and 30 (Figure 5B). Taken together, *ABCB1* was likely the most stable gene in these analyses.

### Comparative analysis of variants between *ABCB1* and *CFTR*

The genomic instability of the coding regions of the *CFTR* gene is of great interest because CFTR mutations are frequent drug targets for therapeutics of cystic fibrosis [reviewed in (Chen et al., 2019)]. Currently, approximately 80% of variants (out of 374 based on a study from 89,052 patients [http://cftr2.org/]), were associated with the development of cystic fibrosis in patients. We performed detailed analyses of sequence motifs of the two ABC transporters using the sequence alignment tools of the UniProt database (www.uniprot.org) (Figure 6). In contrast to the highly conserved *ABCB1*, *CFTR* showed greater variations in the transmembrane domains (TMDs) and the nucleotide binding domains (NBDs) motifs (Figure 6), which are important sites for implementing drug binding, ATP hydrolysis, and drug or ion transport.

**Figure 6.**
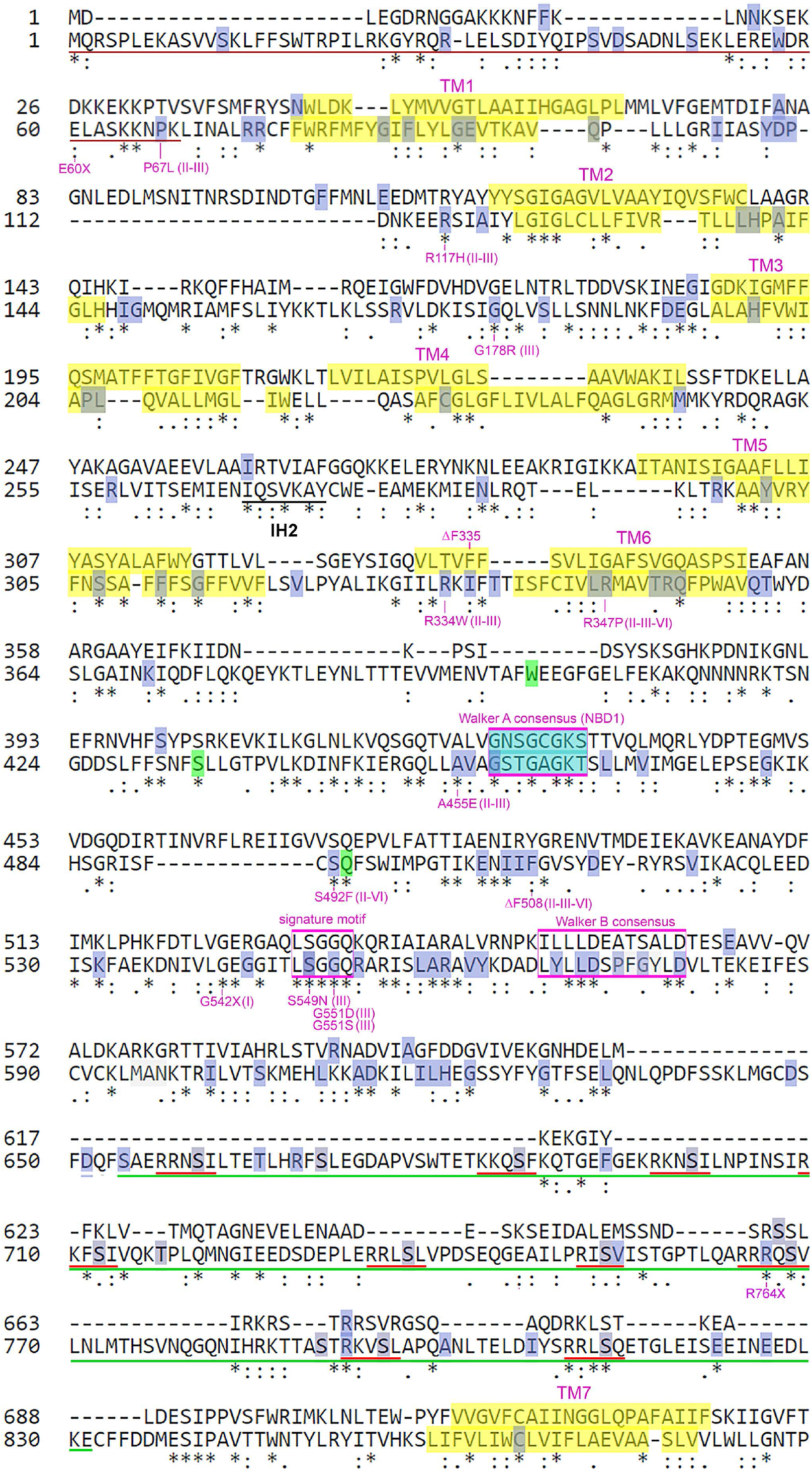
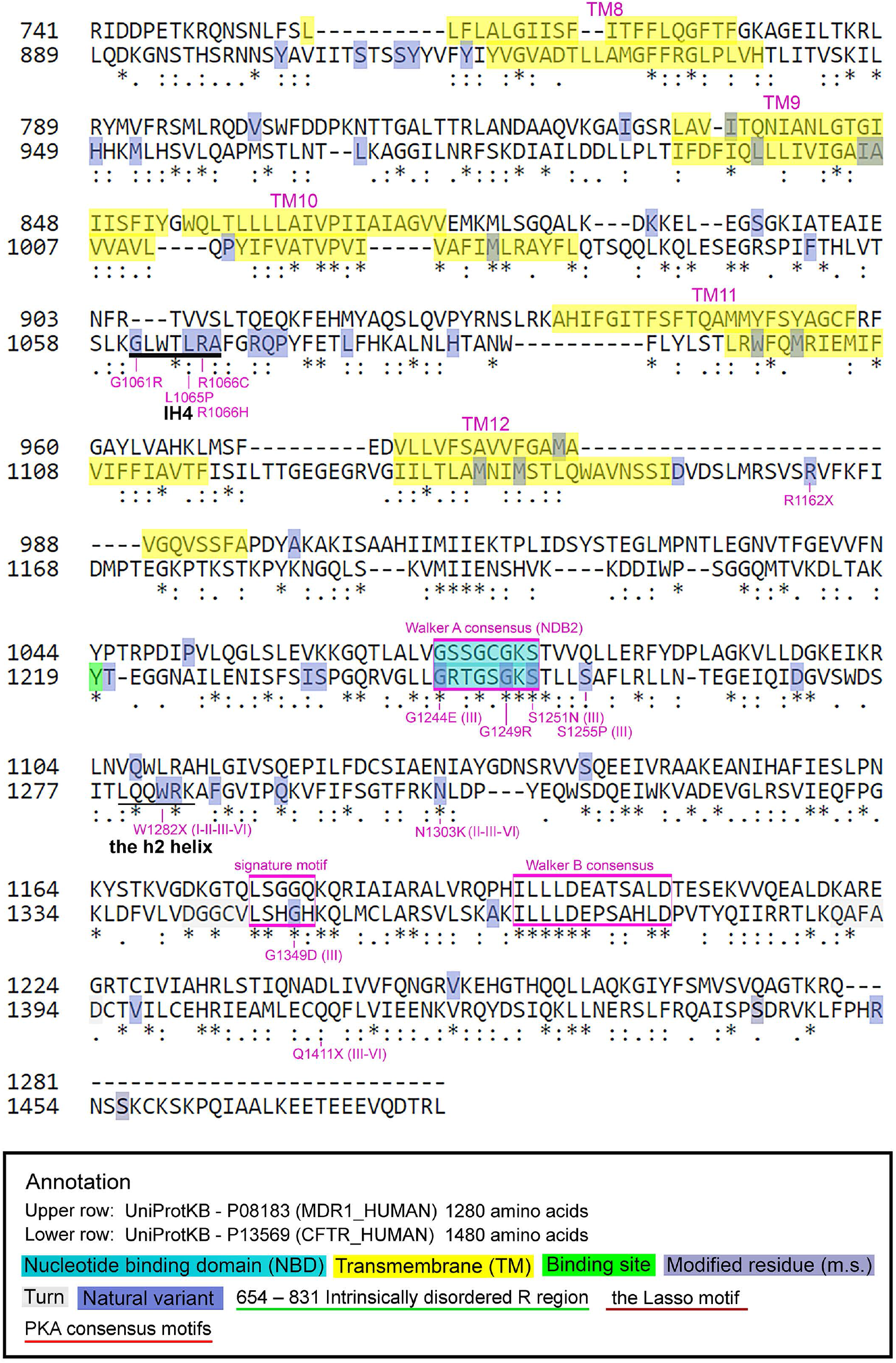
Coding sequence alignments and variant analysis between human ABCB1 and CFTR. The sequence references UniProt P08183 (MDR1/ABCB1) and P13569 (CFTR) were used for the analysis. Nucleotide binding domain (NBD), transmembrane (TMs), modified residues, natural variants, and notable motifs were annotated based on the database information and the analysis. All data were retrieved from UniProt (www.uniprot.org) as of December 2018.

Some representative variants of CFTR include R334W (TM6), A455E (NBD1), F508del (NBD1-TMD interface), G551D (NBD1), G1249R (NBD2), and S1251N (NBD2) (Figure 6). Of note, F508del (the deletion of phenylalanine at the residue 508 of CFTR), is the most common mutation that occurs at a frequency of 70%. Nearly 50% of patients with cystic fibrosis are homozygous for F508del (Cheng et al., 1990; Dalemans et al., 1991; Lukacs et al., 1993; Sharma et al., 2001). With respect to ABCB1, besides our previously reported F335del at the transmembrane 6 (TM6), there are currently no variations or mutations in these corresponding domains in P-gp based on the UniProt database. Thus, this sequence comparison further supported our finding on the genomic stability of the *ABCB1* coding regions.

## DISCUSSION

Concerning the coordinate regulation of both MDR and ADR, there are two outstanding questions to address. One, do *ABCB1* mutations play a major role in conferring the insensitivity to MDR modulation? Second, if not, how do regulatory mechanisms of *ABCB1* facilitate the development of non-*ABCB1*/P-gp mechanisms? The first question is readily answered based on this study: *ABCB1* variations or mutations in the coding regions do not appear to play a significant role in rendering cancer cells resistant to MDR modulation based on our leukemia models and bioinformatics data mining. To address the second question, we propose a hypothesis that coordinates *ABCB1* mutational events with the activation of *ABCB1* and the induction/selection of ADR.

As illustrated in Figure 7, we propose an ADR coordination model, in which the low to no mutational events of the *ABCB1* coding regions lead to expression or overexpression of *ABCB1* mRNAs by stabilizing the native *ABCB1* promoter (designated as P1) or the upstream *ABCB1* promoter (P2) (Figure 4), and by selection of various gene rearrangements at the upstream regions (Figure 7, A1) (Chen et al., 2005; Huff et al., 2005; Mickley et al., 1998; Patch et al., 2015). The stably expressed wild-type *ABCB1* transcripts as well as P-gp may offer cancer cells a mechanism that compensates the dominant effects of a potential mutant *ABCB1*/P-gp on cytotoxic insults. Thus, the expressed wild-type P-gp is partially inhibitable by weaker and non-specific P-gp inhibitors (e.g., CsA) and completely inhibitable by potent and specific P-gp inhibitors (e.g., PSC). Such differential inhibition of P-gp function would conceivably lead to two different trajectories for the development of non-P-gp ADR (Figure 7A, 4-6).

**Figure 7.**
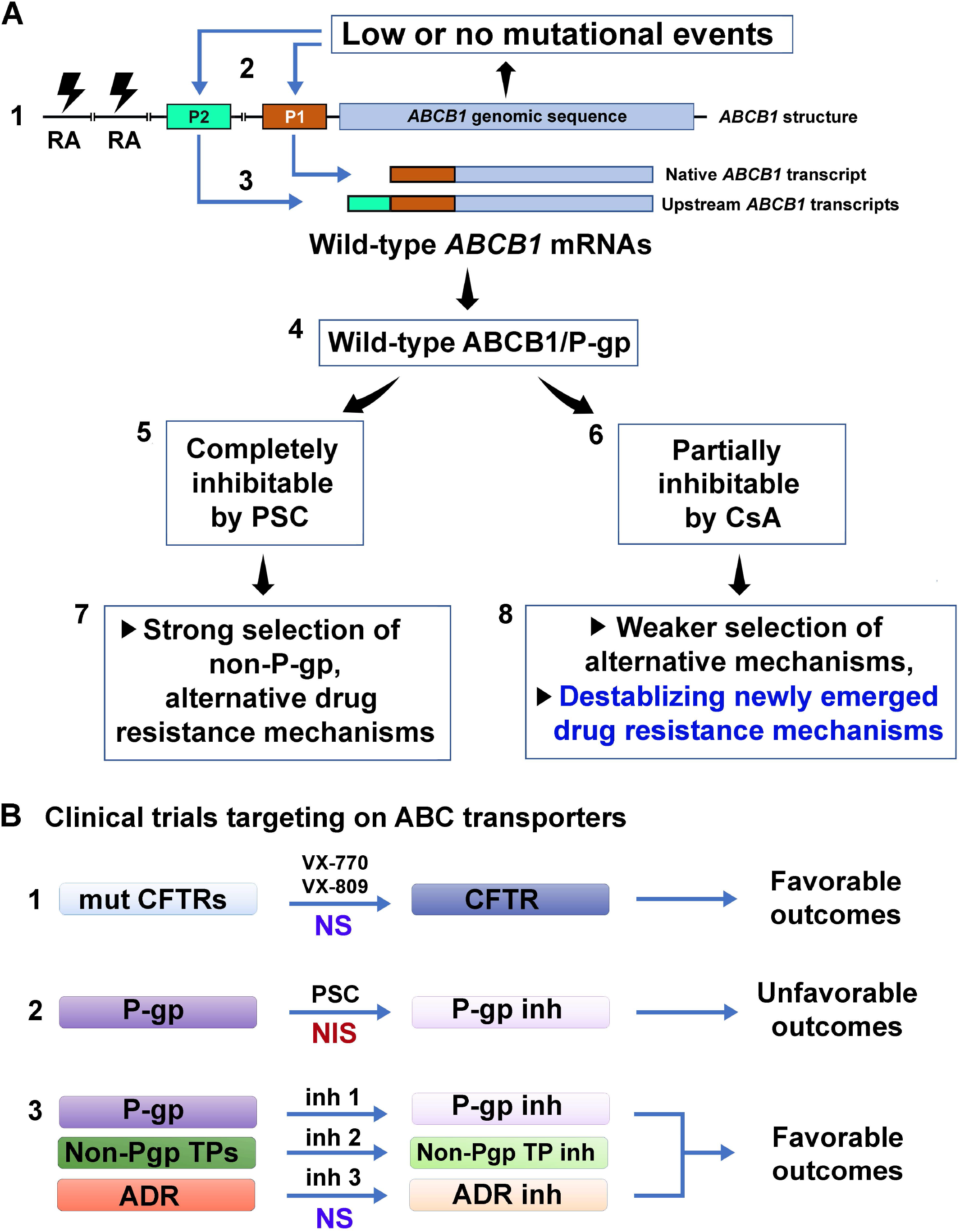
Major pathways that lead to the development of alternative drug resistance mechanisms in human cancers. (**A**) The model is based on the evidence from K562/R7, RVC, RDC, and other drug-resistant cell lines (described in Tables 1 and 2). The model depicts that the low to no mutational events of the *ABCB1* coding regions lead to expression or overexpression of wild-type ABCB1/P-glycoprotein, whose functional inhibition and partial inhibition encourages the development of stable and unstable alternative drug-resistant mechanisms, respectively. (**B**). Therapeutic strategies related to clinical trials that target ABC transporters. Abbreviations: ADR, alternative drug resistance mechanism(s); CsA, cyclosporine; GR, gene rearrangements; Inh, inhibitor(s) or inhibition; mut, mutant(s), NIS, a necessary but insufficient strategy; NS, a necessary and sufficient strategy; P1, the native *ABCB1* promoter located at exon 1 of the gene; P2, the far upstream *ABCB1* promoter located at exon −1 of the gene; P-gp, P-glycoprotein; PSC, SDZ PSC-833 known as valspodar; TPs, transporters; VX-770, the potentiator drug VX-770 (known as ivacaftor) that targets the CFTR-G551D mutation; VX-809, the corrector drug VX-809 (lumacaftor) that targets the CFTR-F508del mutation.

In the case of CsA, it partially inhibits the function of P-gp as well as other multidrug transporters such as ABCG2 and ABCC1 (Qadir et al., 2005). The benefit of such a broad and partial inhibition of diverse multidrug transporters would make cancer cells moderately resistant to anticancer drugs, and much less permissive to the selection of non-transporter-mediated ADR. The CsA-mediated inhibition might also destabilize the newly emerged drug resistance mechanisms (Figures 1, 7A, and Table 1), which may be partially associated with the capacity of CsA to induce apoptosis in some human cancer cells (Sato et al., 2011). However, the disadvantage of such inhibition would be that it could not eradicate MDR cells. Overall, CsA contributes to deferring stable drug-resistant phenotypes in RVC, RDC, and KCVB2 cell lines, which at least partially explains the benefits of CsA in reversing clinical MDR in P-gp-expressing leukemia patients (List et al., 2001).

In the case of PSC, which represents a potent and more specific P-gp inhibitor, it greatly inhibits the function of P-gp as well as a few multidrug transporters such as ABCG2 and ABCC1 (Chen and Sikic, 2012; Robey et al., 2018). A strong inhibition of a wider range of multidrug transporters would sensitize cancer cells to anticancer drugs that are typical P-gp substrates. However, such inhibitory effects would also permit the selection or induction of non-ABC transporter-mediated ADR. This might explain the selection of relatively stable drug-resistant phenotypes in KCVB2, KPTA5, and MES-SA/DxP cell lines using PSC rather than CsA (Figure 1; Figure 7A, 5-6) (Chen et al., 1997; Dumontet et al., 2004; Jaffrezou et al., 1995). The emergence of co-existence of alternate drug resistance mechanisms is likely to explain the unfavorable outcomes identified in the clinical trial that used PSC and other potent P-gp inhibitors as modulators of MDR.

Whether the inhibition of *ABCB1*/P-gp or other ABC drug transporters is a rational therapeutic approach to circumventing clinical MDR is still elusive because of no conclusive clinical data. However, we may address this complicated question by asking a simple one: is targeting the ABC transporter a rational strategy to treat ABC transporter-related diseases? The answer for the latter question is yes. As exemplified by CFTR, a non-drug ABC transporter that functions as a chloride channel, its dominant mutations are now widely used as molecular targets for therapy of patients with cystic fibrosis (CF).

The recent proof-of-concept experiments using stem-cell organoids from patients with CF confirmed that targeting CFTR, an ABC-based chlorine channel, with small molecules is both rational and effective (Chen et al., 2019; Dekkers et al., 2016). Both genetically engineered CFTR cells lines (e.g., transfected diverse CFTR mutant cDNAs) and stemcell organoids from patients with CF can be used to analyze wild-type and mutant CFTR activity *in vitro*, predict drug response for CF patients, and guide personalized drug screens and therapy (Chen et al., 2019).

However, in contrast to CF patients, which is principally associated with the only ABC transporter CFTR, patients with refractory cancer usually express several ABC transporter-dependent and -independent mechanisms. Thus, targeting ABC multidrug transporters for treating patients with MDR can be complicated by the redundancy of these multidrug transporter and other ADR mechanisms. Thus, it is conceivable that Pgp inhibition is still necessary, but insufficient, for reversal of clinical MDR (Figure 7, B2). A combined inhibitory regimen that targets P-gp, non-P-gp transporters, and ADR might be both necessary and sufficient to reverse clinical MDR (Figure 7, B3). Alternatively, MDR may be circumvented clinically by using anticancer agents that are not transport substrates for P-gp and other transporters.

Currently, to evaluate cancer drug response in animal models is difficult as human cancer patients often have considerable genetic heterogeneity, with various non-P-gp genomic mutations and epigenomic states. Cancer stem cell-derived 3D organoids may faithfully retain genetic epigenetic signatures of original cancer tissues [reviewed in (Chen et al., 2018; Sachs and Clevers, 2014)]. Therefore, it is possible that cancer organoids from drug-resistant patient groups might be classified, assayed, and used for predicting drug responders for patients with MDR *in vitro*.

## CONCLUDING REMARKS

Clinical drug resistance is a complicated genetic phenotype, which is attributable to the predominant multidrug transporter ABCB1/P-gp, miscellaneous non-P-gp transporters, and other ADR determinants. Our study in leukemia models revealed that *ABCB1* coding regions are stable even under stringent drug selection, thus suggesting that a mutant Pgp is not a major factor responsible for the unsuccessful modulation of clinical MDR. Partial inhibition of P-gp and other ABC drug transporters by CsA favors induction of unstable drug-resistant phenotypes, which might delay the onset of drug-resistant cells under certain circumstances and benefit some cancer patients. The ADR coordination model supports the rationale for P-gp inhibition as a treatment approach for cancer patients. Co-inhibition of non-P-gp-mediated and ADR mechanisms may be required for overcoming MDR in refractory cancer patients.

## MATERIALS AND METHODS

### Chemical reagents and drugs

The sources of anticancer drugs and chemicals are: etoposide (known as VP-16) was obtained from Bristol-Myers (Evansville, IN), doxorubicin from Adria Laboratories (Columbus, OH), vinblastine from Eli Lilly and Co. (Indianapolis, IN), PSC 833 (known as valspodar) from Sandoz Pharmaceutical Corporation (Basel, Switzerland), cyclosporine (CsA) from Sigma Chemical Co. (St. Louis, MO). and rhodamine 123 (Rh-123) from Molecular Probes (Eugene, OR). All radiolabeled reagents [^3^H]-daunorubicin, [^3^H-vinblastine, and [3H]-etoposide were purchased from Amersham (Arlington Heights, IL). All other chemicals were Sigma-Aldrich (St. Louis, MO).

### Cell culture and development of drug-resistant cellular models

We have previously developed and characterized a panel of drug-resistant cell lines from a human uterine sarcoma, MES-SA and MES-SA/Dx5 (Harker et al., 1983; Harker and Sikic, 1985). Some of which were used as controls in this study (Table 1). In this study, we characterized multiple drug-resistant models from the K562/R7 (R7) cell line by stepwise co-selection of R7 cells with both doxorubicin and CsA, or etoposide and CsA (Figure 1). These cells were cultured in suspension in McCoy’s medium supplemented with in 10% newborn calf serum (Life Technologies, Inc.), 2 mM L-glutamine, and antibiotics (streptomycin and penicillin). Cells were maintained at 37°C in an incubator containing humidified atmosphere and 5% CO_2_. All cultures were routinely tested for mycoplasma infection.

### Drug sensitivity assays

The drug sensitivity or resistance was determined by the MTT [3-(4,5-dimethylthiazol-2-yl)-2,5-diphenyltetrazolium bromide] assay in quadruplicate in 96-well plates, as previously described (Chen et al., 1994).

### Flow cytometric analysis

K562, K562/R7, RVC, and RDC cells were cultured as suspension, counted, filtered through a nylon filter, and centrifuged at 4°C. The cell pellets were resuspended in a modified HBSS buffer, which contained 10 mM 4-(2-hydroxyethyl)-l-piperazineethane sulfonic acid and 5% newborn calf serum. Approximately, 3 x 10^5^ cells were incubated with CsA and PSC at various indicated concentrations (Figure 2) for 1 hour prior to the addition of Rh-123 (0.1 μg/mL) and followed by 45-minute incubation at 37°C. Rh-123 retention was assessed by a laser flow cytometer (FACS-II; Becton-Dickinson Corp., Mountain View, CA).

### [^3^H]-labeled drug accumulation

Intracellular drug accumulation was implemented by adding [^3^H]-daunorubicin (4 μM), [^3^H]-vinblastine (50 nM), and [^3^H]-etoposide (10 mM) to exponentially growing cells. After incubation at 37°C for 1 hour, cells were spun through Nyosil oil to separate them from drug-containing medium. The cell pellets were solubilized in 4% sodium dodecyl sulfate (SDS) solution, incubated at 65°C for 1 hour, and suspended with the Ecolite Liquid Scintillation Cocktail (MP Biomedicals, Inc.). The radioactivity in these samples was measured with a Beckman LS-9000 scintillation counter.

### RNase protection assay

The RNase protection assay, used to examine *ABCB1* mRNA levels and to map the transcription start sites of the *ABCB1* gene, was performed as previously described (Chen et al., 2005; Mickley et al., 1998). Briefly, total RNAs were prepared from parental K562, K562/R7, RVC, and RDC cells. An antisense RNA probe was produced by SP6 RNA polymerase using the 982-bp genomic fragment (of the *ABCB1* proximal promoter) as the template. Total RNAs were hybridized with [^32^P] radiolabeled antisense RNA probes (~2 × 10^5^ cpm).

### Data mining from UniProt

We analyzed the frequency of natural variants of several representative human ABC transporters based on the UniProt database (www.uniprot.org) using the bioinformatic tools of the database in December 2018. ABC Protein entry names are: ABCA1 (O95477, ABCA1_HUMAN, 2261 amino acids), ABCB1 (P08183, MDR1_HUMAN, 1280 amino acids), ABCC1 (P33527, MRP1_HUMAN, 1531 amino acids), ABCC7/CFTR (P13569, CFTR_HUMAN, 1485 amino acids), and ABCG2 (Q9UNQ0, ABCG2_HUMAN, 655 amino acids).

## ACKNOWLEDGMENTS

This work was supported by National Cancer Institute grant CA09302 (K.G.C), NIH grants R01 CA52168, R01 CA92474, and R01 CA114037 (BIS), American Cancer Society Grant DHP-76E (B.I.S.), and in part by the Intramural Research Program of the NIH at the National Institute of Neurological Disorders and Stroke (K.G.C.). We thank Dr. William H. Fleming for technical assistance with flow cytometric experiments and Alicia A. Livinski, NIH Library, for manuscript editing assistance.

